# Selectivity among anti-σ factors by *Mycobacterium tuberculosis* ClpX influences intracellular levels of Extracytoplasmic Function σ factors

**DOI:** 10.1101/491043

**Authors:** Anuja C Joshi, Prabhjot Kaur, Radhika K Nair, Deepti S Lele, Vinay Kumar Nandicoori, Balasubramanian Gopal

## Abstract

Extracytoplasmic Function σ factors that are stress inducible are often sequestered in an inactive complex with a membrane-associated anti-σ factor. *M. tuberculosis* membrane-associated anti-σ factors have a small stable RNA gene A-like degron for targeted proteolysis. Interaction between the unfoldase, ClpX, and the substrate with an accessible degron initiates energy-dependent proteolysis. Four anti-σ factors with a mutation in the degron provided a set of natural substrates to evaluate the influence of the degron on degradation strength in ClpX-substrate processivity. We note that a point mutation in the degron (XXX-Ala-Ala) leads to an order of magnitude difference in the dwell time of the substrate on ClpX. Differences in ClpX/anti-σ interactions were correlated with change in unfoldase activity. GFP chimeras or polypeptides of identical length with the anti-σ degron also demonstrate degron-dependent variation in ClpX activity. We show that degron-dependent ClpX activity leads to differences in anti-σ factor degradation thereby regulating the release of free σ from the σ/anti-σ complex. *M. tuberculosis* ClpX activity thus influences changes in gene expression by modulating the cellular abundance of ECF σ factors.

**Importance:** The ability of *Mycobacterium tuberculosis* to quickly adapt to the changing environmental stimuli occurs by maintaining protein homeostasis. Extra-cytoplasmic function (ECF) σ factors play a significant role in coordinating the transcription profile to changes in environmental conditions. Release of the σ factor from the anti-σ is governed by the ClpXP2P1 assembly. *M. tuberculosis* ECF anti-σ factors have a ssrA-like degron for targeted degradation. A point mutation in the degron leads to differences in ClpX mediated proteolysis and affects the cellular abundance of ECF σ-factors. ClpX activity thus synchronizes changes in gene expression with environmental stimuli affecting *M. tuberculosis* physiology.

## Introduction

*Mycobacterium tuberculosis* encounters diverse host microenvironments including acidification of phagosomes, nitrogen intermediates, reactive oxygen species, nutrient starvation, DNA damage, phosphate deprivation and hypoxia (1). Extracytoplasmic Function (ECF) σ factors are non-essential and stress inducible and they contribute significantly to bacterial survival alongside one- and two-component systems (2). *M. tuberculosis* has ten ECF σ factors-of which four are localized in an inactive complex with membrane associated anti-σ factors (3, 4). The membrane associated anti-σ factors (RsdA, RsmA, RskA and RslA in *M. tuberculosis*) share a common structural organization comprising of an extra-cytoplasmic domain that is a receptor for environmental stress connected to the cytoplasmic anti-σ domain by a single transmembrane helix (Figure 1). The stress-induced release of an ECF σ factor from the σ/anti-σ factor complex governs the intra-cellular levels of these transcription initiation factors and thereby the expression of their cognate regulons. The relative cellular abundance of different σ factors dictates the expression profile-best described by a mechanistic model referred to as the partitioning of σ factor space (5). Indeed, the number of different σ factors is correlated with the diversity of environmental conditions encountered by the bacterium (5).

**Figure 1:**
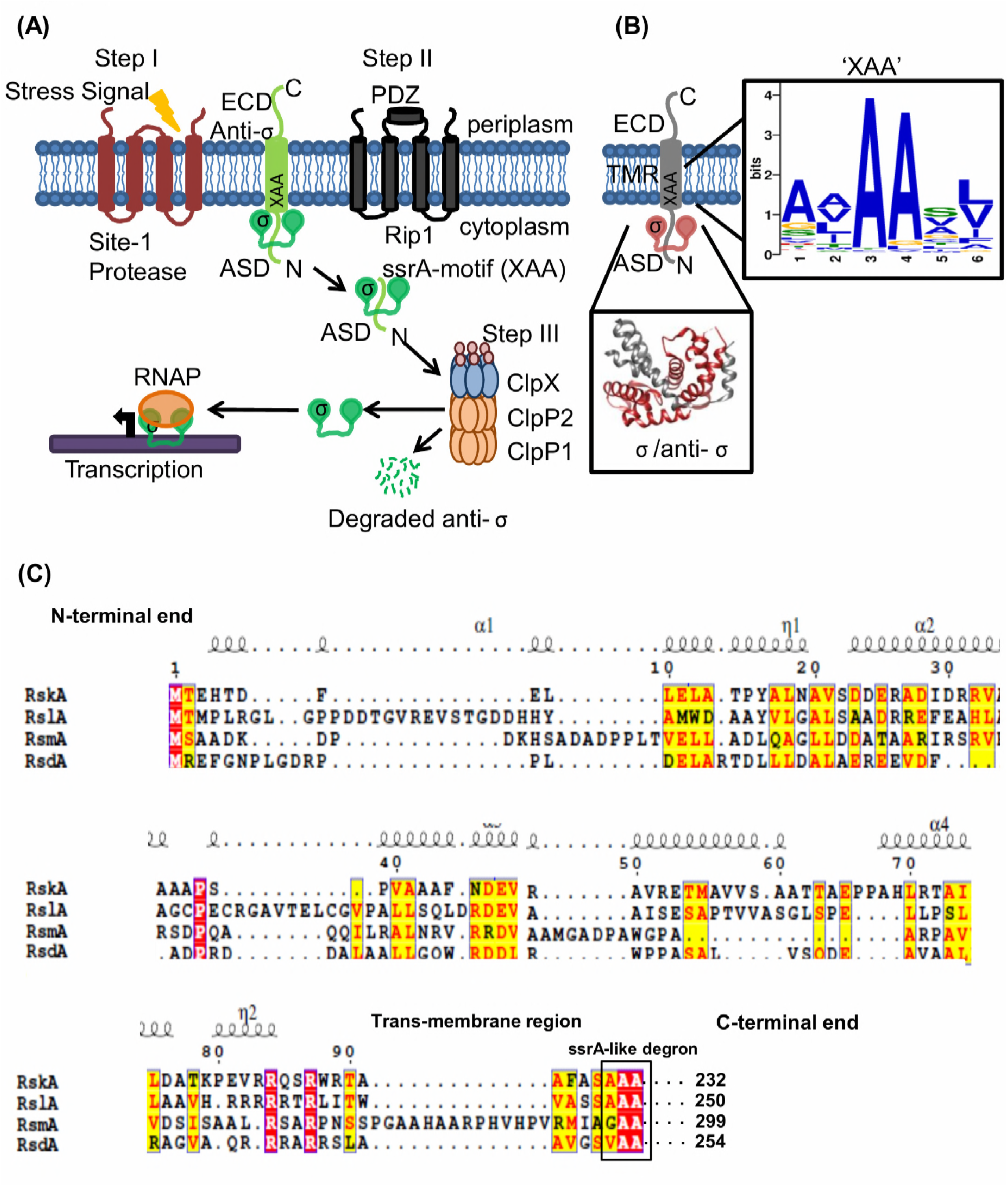
Regulated Intra-membrane proteolysis in *M. tuberculosis.* **(A)** Schematic of the Regulated Intra-membrane Proteolysis (RIP) pathway in *M. tuberculosis*. The site-1 protease that has not been identified thus far (step I) triggers the proteolytic cascade by cleaving the Extra-cytoplasmic domain (ECD) of the anti-σ factor. After the activity of the site-2 protease, Rip1, the ssrA-like motif is exposed and intracellular proteolysis (by the ClpXP2P1complex, step III) occurs by selectively degrading the anti-σ domain (ASD), governing the cellular abundance of Extra Cytoplasmic Function (ECF) σ factors. **(B)** All ECF anti σ-factors with one trans-membrane helix have a ssrA-like degron in the trans-membrane region (TMR). The anti σ-factors used for this analysis were from the publised curated dataset (63). The trans-membrane region was identified using the TMHMM server, version 2.0 (70). The degron (inset) is highlighted using the MEME suite (71). **(C)** Sequence features of the four anti-σ domains (cytosolic fragment) of the membrane-associated *M. tuberculosis* anti-σ factors (aligned using ESpript 3.0) (72). The degron at the C-terminus of the cytosolic domain is made accessible after Rip1 proteolysis (step II); Rip1 activity dissociates the σ/anti-σ complex from the membrane.

The intracellular release of an ECF σ factor from the inactive membrane-associated σ/anti-σ complex is governed by a proteolytic cascade referred to as the Regulated Intra-membrane Proteolysis (RIP) pathway (6). This cascade is initiated by the action of a so-called site-1 protease that acts on the extracytoplasmic domain of the anti-σ factor (6). This triggers the activity of a trans-membrane protease (site-2 protease) that dissociates the σ/anti-σ complex from the membrane. The anti-σ factor is then degraded by energy-dependent proteolytic complexes to release the bound ECF σ factor that can associate with the RNA polymerase and initiate transcription (Figure 1A). The intracellular proteolysis of the anti-σ factor RseA is primarily governed by ClpXP in *Escherichia coli,* although other proteolytic assemblies also contribute to this process (7). The specific degradation of *E. coli* RseA from the σ^E^/RseA complex is also influenced by an adaptor protein, SspB (8). *E. coli* SspB mediated interactions are crucial for effective degron recognition-while *E. coli* ClpX interacts with residues 9-11 at the C terminus of the ssrA degron, SspB interacts with residues 1-4 and 7 (9-11). Other *E. coli* ClpX adaptors that have been characterized are RssB and UmuD (12,13). The presence of different adaptors suggested a mechanism for the specific recruitment of diverse substrates for the ClpX unfoldase to initiate proteolysis with the serine protease ClpP in the ClpXP proteolytic complex (12).

ClpX comprises of a small N-terminal domain flexibly attached to the unfoldase module, the AAA+ domain (14). The AAA+ domain has multiple conserved sequence features including Walker A and Walker B motifs for ATP binding, a second region of homology (SRH) segment involved in ATP hydrolysis and sensor 2 and 3 residues that propagate conformational changes upon ATP hydrolysis to stabilize the ATP binding conformation of the unfoldase (14-18). With the two domains functioning in a concerted manner, ClpX can translocate and unfold a diverse range of substrates (19). Analysis of *E. coli* ClpX substrates suggested five distinct degron motifs (19). Apart from adaptor proteins that enforce specificity, the N-terminal domain of *E. coli* ClpX is also involved in substrate recognition (15). The role of the ClpX N-terminal domain, however, differs across substrates. While the N-terminal domain substantially influences *E. coli* ClpX action on substrates like λO and MuA, it is much less so for Green Fluorescent Protein (GFP) substrates with a small stable RNA gene A (ssrA) degron (15).

The *M. tuberculosis* RIP pathway is only partially characterised. The site-1 protease that initiates the proteolytic cascade in the RIP pathway has not been identified in *M. tuberculosis.* One site-2 protease Rip1 (Rv2869c) acts on all membrane associated anti-σ factors (RskA, RsmA, RslA and RsdA) (20,21). For comparison, in *E. coli,* the first two proteolytic steps are performed by DegS and YaeL (22,23). However, straightforward extension from the *E. coli* model for the subsequent steps is difficult as there are four membrane-associated σ/anti-σ complexes in *M. tuberculosis* (σ^D^/RsdA, σ^K^/RskA, σ^L^/RslA and σ^M^/RsmA) as opposed to one (σ^E^/RseA) in *E. coli* (Figure 2A). The cytosolic step of the cascade involving intracellular proteolytic complexes is also substantially different in *M. tuberculosis* than either *E. coli* or *B. subtilis* (8,24). For example, *M. tuberculosis* has two ClpP protease components-ClpP1 and ClpP2 (25). Furthermore, targeted protein degradation in *E. coli* by the ClpXP complex is also influenced by adaptor proteins. Unlike *E. coli*, no SspB homologue or adaptor of *M. tuberculosis* ClpX has been annotated or experimentally identified thus far. Nonetheless, previous studies revealed that the cytoplasmic domain of RsdA was recognised and cleaved by the *M. tuberculosis* ClpXP2P1 complex (26). The degron in *M. tuberculosis* RsdA is VAA, identical to that in *E. coli* RseA. RslA, however, was found to be resistant to ClpXP2P1 degradation, despite having the ssrA-like degron (26). Apart from proteolytic degradation, other mechanism(s) modulate the cellular abundance of specific ECF σ factors by altering the rates of an ECF σ from an inactive σ/anti-σ complex. For example, *M. tuberculosis* RslA was shown to release σ^L^ under oxidative stress conditions (27). In this case, the receptor for the redox stimulus was the Zinc binding CXXC motif in the anti-σ factor, RslA. The release of Zinc under oxidizing conditions was seen to alter the conformation of RslA thereby releasing σ^L^. *M. tuberculosis* RskA also was shown to dissociate under reducing conditions from σ^K^, the redox sensor in this case, is the σ factor σ^K^ (28). All these anti-σ factors, however, also contain an ssrA-like degron that is exposed upon RIP-1 (site-2 protease) activity.

**Figure 2:**
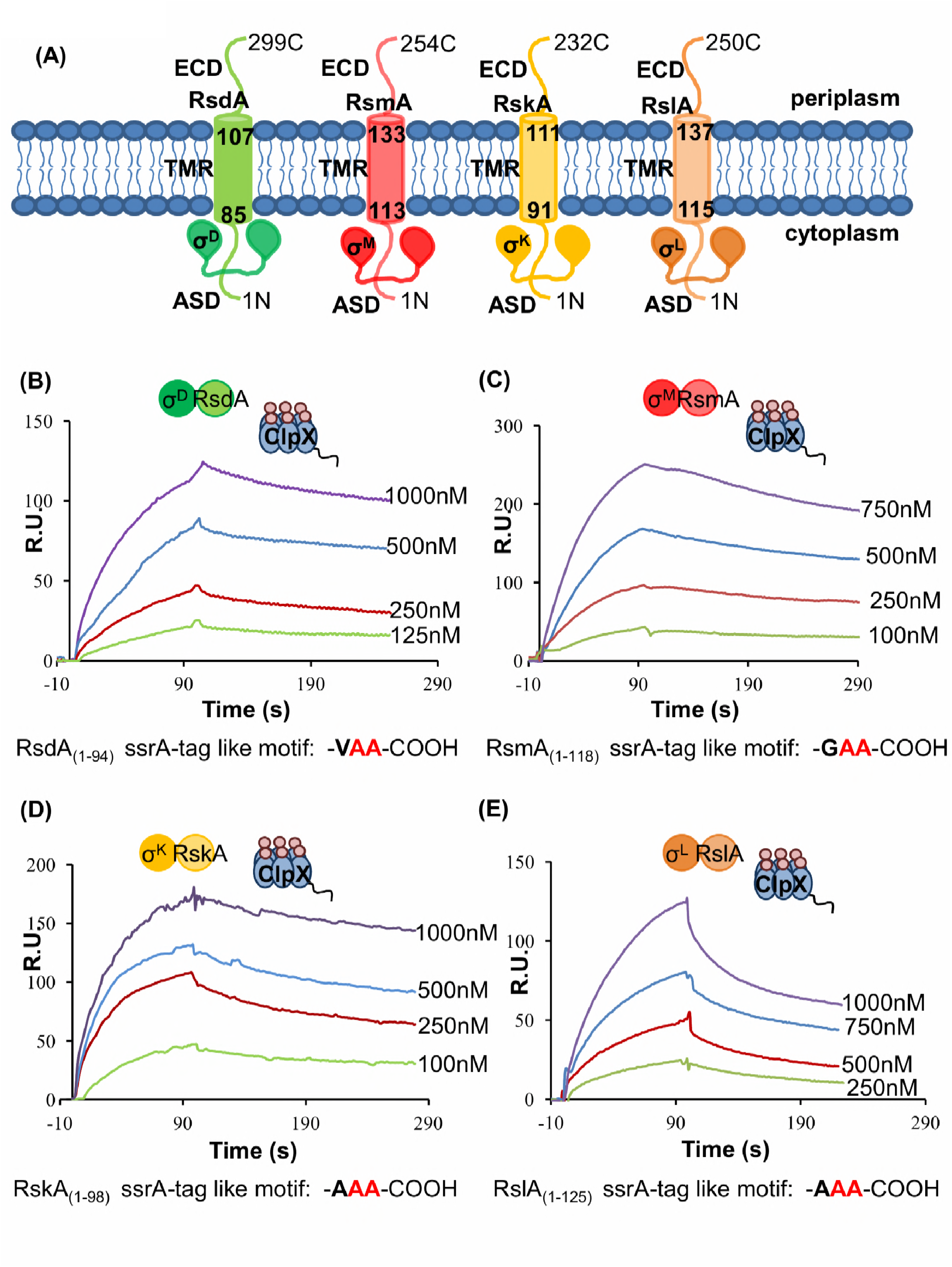
SPR sensorgrams of ClpX interactions with the four σ/anti-σ complexes. **(A)** Schematic representation of the *M. tuberculosis* σ factors in complex with the membrane anchored anti-σ factors. **(B)** SPR sensorgram of σ^D^/RsdA-ClpX interactions. σ^D^/RsdA binds the tightest to ClpX with an affinity *ca* 14.5nM. **(C)** A ten-fold reduction in the binding affinity was seen in the binding of σ^M^/RsmA with ClpX (corresponding to a K_D_ of 1.33*10^−7^ M). **(D)** The binding of σ^K^/RskA with ClpX is intermediate between σ^M^/RsmA and σ^L^/RslA *(ca* 164nM). **(E)** Interaction of σ^L^/RslA with ClpX reveals substantial reduction in the binding affinity when compared to σ^D^/RsdA. The interaction data is compiled in Table 1. Details of the protein constructs are described in supporting information Table A4.

In the regulated proteolytic cascade, the targeted proteolysis of an anti-σ factor by the ClpXP proteolytic complex is the last step in signal transduction to effect changes in gene expression in response to environmental stress. The cytosolic domains of four *M. tuberculosis* anti-σ factors with the ssrA-like degron provided a set of natural variants to understand the basis for substrate selection in *M. tuberculosis* ClpX. We note that while the N-terminal domain of ClpX is not involved in degron recognition, it influences unfoldase activity. We also describe biochemical experiments which reveal that the degron sequence governs both the substrate binding affinity as well as the kinetics of unfolding. The variation in the dwell time of the substrate on ClpX was also seen to have a direct bearing on the proteolytic degradation of the anti-σ substrates by the ClpXP2P1 complex to release free ECF σ factors that can initiate transcription. In effect, *M. tuberculosis* ClpX translates variation in the degron sequence into differential unfoldase activity. These degron-dependent differences in last step in the *M. tuberculosis* RIP cascade are thus likely to provide an additional regulatory layer for nuanced changes in the transcriptional profile in response to a stress stimulus.

## Results

### The ssrA-like degron governs interactions between *M. tuberculosis* ClpX and anti-σ substrates

The binding affinity of *M. tuberculosis* ClpX with the anti-σ substrates containing the C-terminal ssrA-like degron was determined by Surface Plasmon Resonance (SPR) (Figure 2B-D). As the purified anti-σ factors are prone to aggregation, the purified substrates used in these experiments consisted of the ECF σ factor complexed with an anti-σ factor containing the degron at the C-terminus. Co-expression and co-purification of the σ/anti-σ factor complexes significantly improves the yield of homogenous protein samples as the σ or anti-σ factors in isolation are relatively unstable (29). The last three residues (9-11) of the ssrA degron was shown to interact with *E. coli* ClpX (19). This feature was seen to be retained in the case of *M. tuberculosis* ClpX (26). SPR sensorgrams revealed that deletion of the terminal residues of the degron abrogated the binding of these substrate proteins to ClpX-a finding that was similar to the case where the last three residues of the degron (VAA) were replaced by a negatively charged C-terminus (VDD) (Figure 3C, Figure A1). Indeed, the finding that substrates without the degron do not bind ClpX also suggests that non-specific binding is unlikely (Figure A1). These constructs were employed as control inactive ‘degrons’ in the subsequent analysis. *M. tuberculosis* ClpX bound to σ^D^/RsdA with the highest affinity (Table 1). We note a ten-fold reduction in ClpX binding to the σ^L^/RslA, σ^K^/RskA and σ^M^/RsmA complexes as compared with the the σ^D^/RsdA substrate (Table 1). These SPR measurements were performed using freshly prepared ClpX samples. We note that ATP does not appear to be a necessary prerequisite for substrate binding. This differs from previous reports that suggested nucleotide addition as a trigger for substrate recruitment in this class of unfoldases. A related observation is that while freshly prepared ClpX samples are primarily hexameric with a small dimeric component, the hexameric species is unstable in the absence of ATP (polydispersity increases after *ca* 24 hrs). SPR sensorgrams reveal that the difference in these ClpX-σ/anti-σ interactions lies in the dissociation rate constant (kd) (Table 1). A comparison between the standard deviation in these measurements across different substrates is shown in Figure A2. The degrons in the four anti-σ factors have different aliphatic residues at the ante-penultimate position (Figure 1C). A mutation in the degron of RsdA where the degron resembled that of RslA (AAA) and RsmA (GAA) shows that the binding affinity of ClpX with the σ^D^/RsdA_AAA_ and the σ^D^/RsdA_GAA_ mutant is less than wild-type σ^D^/RsdA - comparable to ClpX interactions with the σ^L^/RslA and the σ^M^/RsmA complexes respectively (Figure 3A-B, Table 2). These observations suggest that although the C-terminal Alanines in the degron are involved in ClpX interactions, the ante-penultimate residue influences substrate tethering and consequently the residence time of the substrate protein on the ClpX hexamer (Table 2).

**Figure 3:**
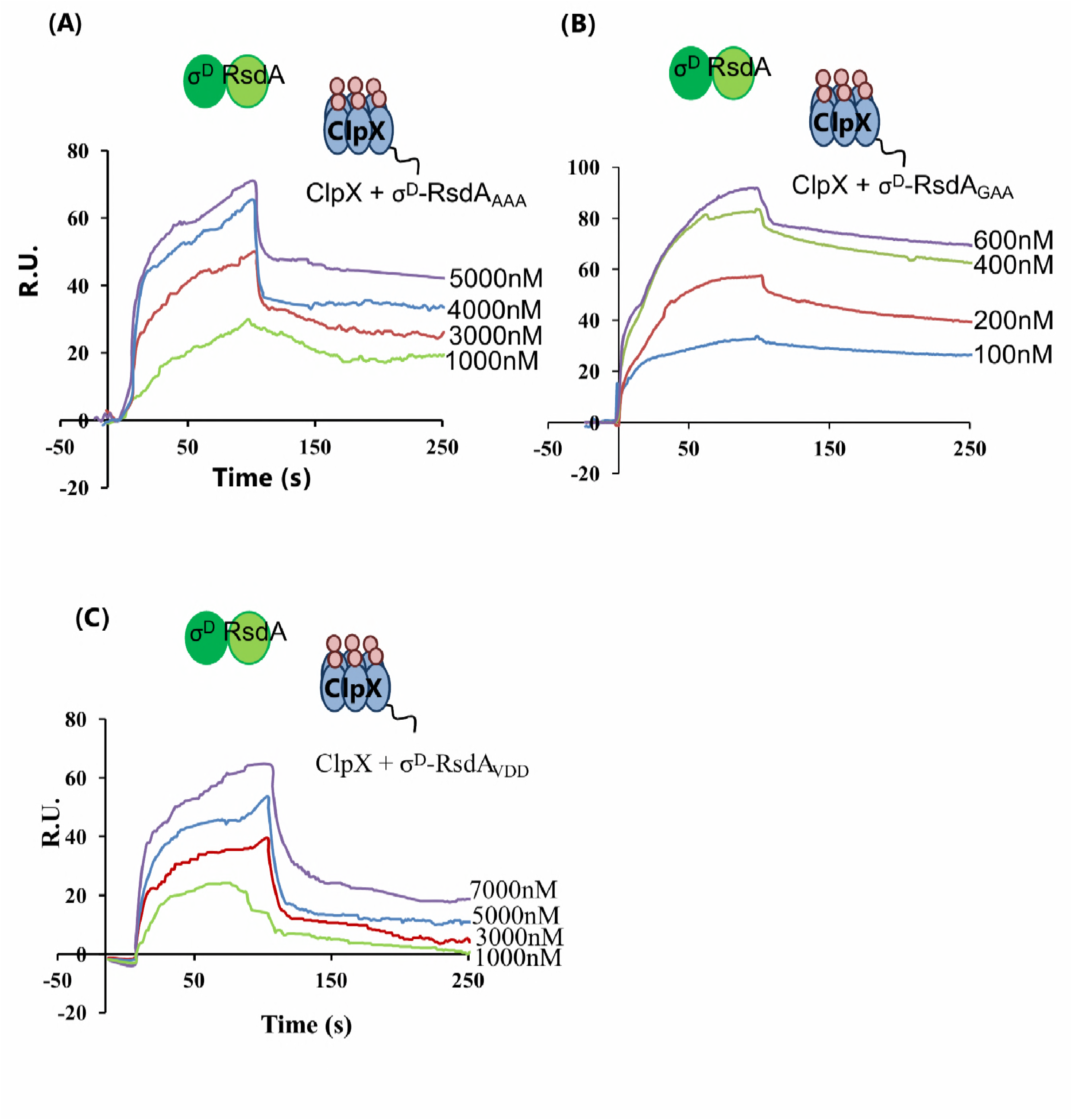
Sequence and conformational features that influence ClpX substrate recruitment in the absence of adaptor proteins. **(A-B)** SPR measurements of σ^D^/RsdA_AAA_ −ClpX and σ^D^/RsdA_GAA_ –ClpX interactions reveal that σ^D^/RsdAAAA and σ^D^/RsdAGAA bind to ClpX with ten-fold lower affinity when compared to the σ^D^/RsdAVAA complex. The binding affinity of the σ^D^/RsdAAAA complex is equal to the σ^L^/RslA complex. **(C)** Charged residues at the terminal end of the degron substantially reduce ClpX-substrate interaction. The affinity of the σ^D^/RsdA_VDD_ complex is hundred-fold lower than that for the σ^D^/RsdA_VAA_ complex.

**Table 1:** Kinetic parameters of ClpX interactions with anti -σ factors.

| **$\sigma$/anti-σ complex** | **Association**<br>**rate constant**<br>**$(k_a)$ ($M^{-1}s^{-1}$)** | **Dissociation**<br>**rate constant**<br>**$(k_d)$ ($s^{-1}$)** | **$K_D$ (M)** | **$\Delta G$ (kJ/mol)** |
| --- | --- | --- | --- | --- |
| $\sigma^D$-RsdA<sub>(1-94)</sub> | $6.4 \cdot 10^4$ | $9.2 \cdot 10^{-4}$ | $2.8 \cdot 10^{-8}$ | -44.66 |
| $\sigma^M$-RsmA<sub>(1-118)</sub> | $2 \cdot 10^4$ | $2.7 \cdot 10^{-3}$ | $1.2 \cdot 10^{-7}$ | -39.22 |
| $\sigma^K$-RskA<sub>(1-98)</sub> | $3.3 \cdot 10^4$ | $5.5 \cdot 10^{-3}$ | $2.2 \cdot 10^{-7}$ | -38.69 |
| $\sigma^L$-RslA<sub>(1-125)</sub> | $3.0 \cdot 10^4$ | $4.9 \cdot 10^{-3}$ | $2.3 \cdot 10^{-7}$ | -38.02 |

**Table 2:** Interactions between σ^D^-RsdA degron variants and ClpX.

| **$\sigma$/anti-σ complex** | **Association rate**<br>**constant**<br>**$(k_a)$ ($M^{-1}s^{-1}$)** | **Dissociation rate**<br>**constant**<br>**$(k_d)$ ($s^{-1}$)** | **$K_D$ (M)** | **$\Delta G$ (kJ/mol)** |
| --- | --- | --- | --- | --- |
| $\sigma^D$-RsdA<sub>VAA</sub> | $6.4 \cdot 10^4$ | $9.2 \cdot 10^{-4}$ | $1.4 \cdot 10^{-8}$ | -44.66 |
| $\sigma^D$-RsdA<sub>GAA</sub> | $4.3 \cdot 10^4$ | $5.8 \cdot 10^{-3}$ | $1.4 \cdot 10^{-7}$ | -35.84 |
| $\sigma^D$-RsdA<sub>AAA</sub> | $6.8 \cdot 10^3$ | $2.7 \cdot 10^{-3}$ | $6.9 \cdot 10^{-7}$ | -36.49 |
| $\sigma^D$-RsdA<sub>VDD</sub> | $1.3 \cdot 10^4$ | $1.3 \cdot 10^{-2}$ | $1.1 \cdot 10^{-6}$ | -33.37 |

### The degron in substrates determines ClpX ATPase and unfoldase activity

Processing of protein substrates by ClpX is coupled to the rate of ATP hydrolysis (30). ATPase assays revealed that differences in binding affinity of the four anti-σ substrates to ClpX were correlated with the rate of ATP hydrolysis. That substrate binding directly influences ATPase activity was evident from the observation that the ATPase activity of ClpX was highest in the presence of σ^D^/RsdA-almost five-fold higher when compared to ClpX alone. Secondly, ClpX ATPase activity was not altered in the presence of the substrate with the inactive degron in σ^D^/RsdA (VDD). The ATPase activity of ClpX in the presence of σ^L^/RslA was three-fold lower than that of σ^D^/RsdA (Figure 4A, Table 3). Given that the lengths of the anti-σ factors vary from 222 to 375 residues, due to the linker between the transmembrane helix and the cytosolic anti-σ domain (Figure 1C, Table A5), the ATPase activity of ClpX was evaluated using one substrate (σ^D^/RsdA) with varying ante-penultimate residues in the degron (Figure 4A). When the antepenultimate residue was modified to Ala/Gly from Val in σ^D^/RsdA, ATPase activity decreased – consistent with SPR experiments that show that ClpX binds this substrate protein with reduced affinity (Figure 4A, Table 3). In another experiment to evaluate the effect of variation of substrate size on the binding affinity or ATPase activity, ClpX ATPase activity was also evaluated in the presence of ssrA peptide chimeras. As seen in Figure 4b, the ATPase activity of ClpX was substantially enhanced in ssrAVAA (mimicking σ^D^/RsdA) but was less affected by ssrA_AAA_ (mimicking σ^L^/RslA) (Figure 4B, Table 3). Finally, we performed an experiment wherein Green Fluorescent Protein (GFP) chimeras with different degrons at the C-terminus were subjected to unfolding by ClpX. The degrons in these GFP-chimera substrates mimic those in the anti-σ factors. These results show that GFP_VAA_ (mimicking σ^D^/RsdA) unfolded faster than GFPGAA (mimicking σ^M^/RsmA) or GFPAAA (mimicking σ^K^/RskA) (Figure 4C, Table A1). As in the case of the σ/anti-σ substrates, GFP_1-8ssrA_ (devoid of the last three residues in the degron) was not unfolded by ClpX. Taken together, the data suggests that degron composition, in particular the ante-penultimate residue, affects the ATPase and unfoldase activity of *M. tuberculosis* ClpX (Figure 4A-C).

**Figure 4:**
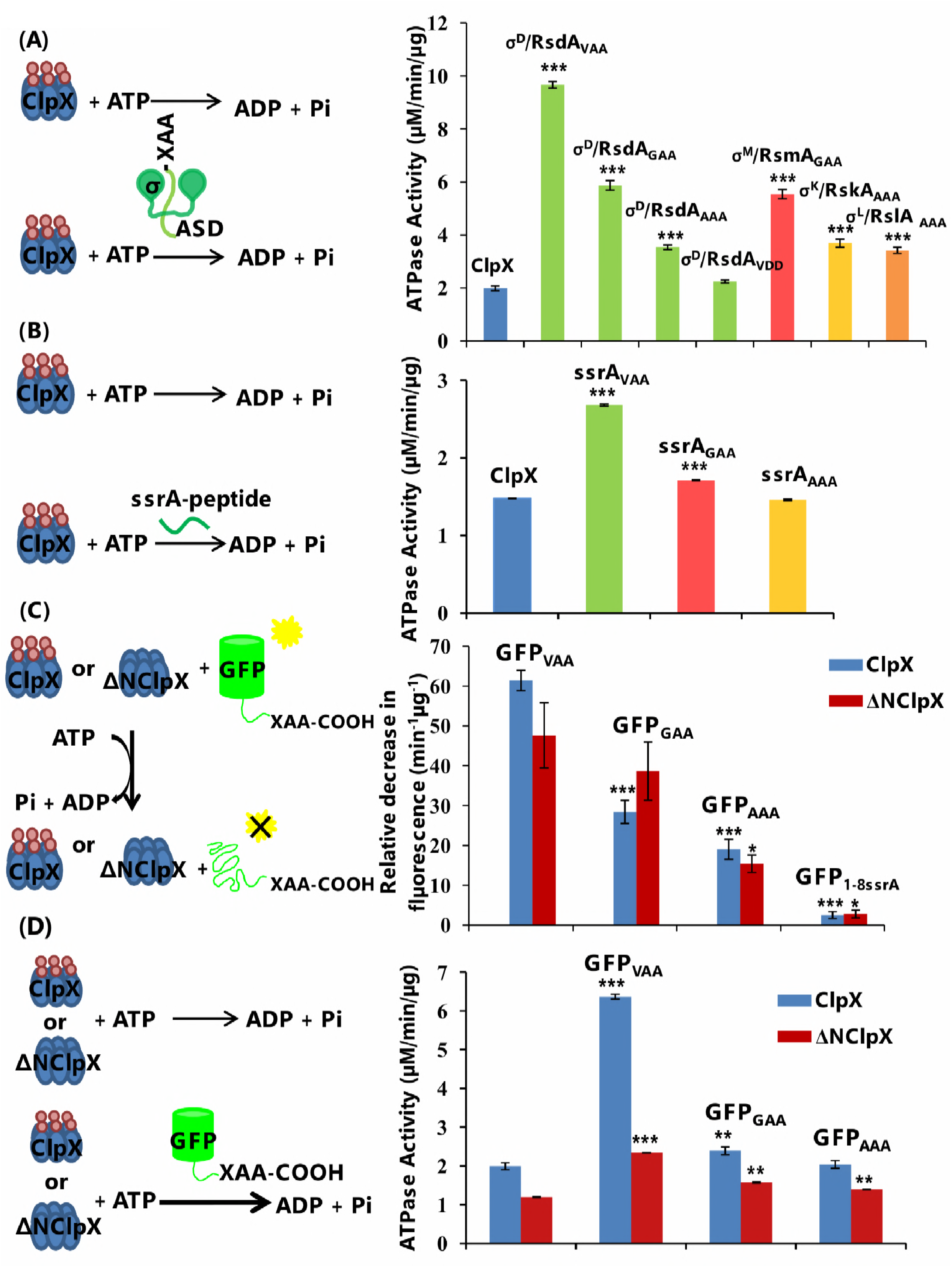
The degron influences the unfoldase activity of *M. tuberculosis* ClpX. **(A)** The ATPase activity of ClpX is altered by the presence of substrates-with maximum activity for the σ^D^/RsdA_VAA_ substrate and the lowest for the σ^D^/RsdAVDD mutant. **(B)** The degron attached to polypeptides of equal length also shows degron-dependent gradation in ATPase activity. ATPase activity of ClpX increases only in the presence of the degron peptide ending with ‘VAA’. No change in ATPase activity was observed for degrons with either ‘GAA’ or ‘AAA’. **(C)** GFP chimeras mimic the anti-σ substrates in inducing degron dependent gradation in unfoldase activity. The relative decrease in fluorescence was highest for GFP_VAA_ followed by GFP_GAA_ and GFP_AAA_ for both full-length ClpX and ΔNClpX. Details of protein constructs are compiled in supporting information Table A4. **(D)** Removal of the ClpX N-terminal domain reduces degron-dependent change in ClpX activity. ** indicates significance at p<0.005, *** indicates significance at p<0.0001.

**Table 3:** ATPase activity of ClpX in the presence of substrates.

| **anti-σ substrates** | **Specific Activity ($\mu\text{M}/\text{min}/\mu\text{g}$)** |
| --- | --- |
| ClpX | $1.9 \pm 0.1$ |
| ClpX+$\sigma^D$/RsdA<sub>VAA</sub> | $9.6 \pm 0.1$ |
| ClpX+$\sigma^M$/RsmA<sub>GAA</sub> | $5.5 \pm 0.2$ |
| ClpX+$\sigma^K$/RskA<sub>AAA</sub> | $3.7 \pm 0.2$ |
| ClpX+$\sigma^L$/RslA<sub>AAA</sub> | $3.4 \pm 0.1$ |
| **RsdA-degtron variants** |  |
| ClpX+$\sigma^D$/RsdA<sub>GAA</sub> | $5.9 \pm 0.2$ |
| ClpX+$\sigma^D$/RsdA<sub>AAA</sub> | $3.5 \pm 0.1$ |
| ClpX+$\sigma^D$/RsdA<sub>VDD</sub> | $2.2 \pm 0.1$ |
| **GFP chimeras** |  |
| ClpX | $1.4 \pm 0.1$ |
| ClpX+GFP<sub>VAA</sub> | $6.4 \pm 0.1$ |
| ClpX+GFP<sub>GAA</sub> | $2.4 \pm 0.1$ |
| ClpX+GFP<sub>AAA</sub> | $2.0 \pm 0.1$ |
| **ssrA peptide** |  |
| ClpX | 1.5 |
| ClpX+ssrA<sub>VAA</sub> | 2.7 |
| ClpX+ssrA<sub>GAA</sub> | 1.7 |
| ClpX+ ssrA<sub>AAA</sub> | 1.5 |

### Role of the N-terminal domain of *M. tuberculosis* ClpX in substrate recruitment

Next, we evaluated the role of the N-terminal domain in stabilizing ClpX-substrate interactions. Towards this ATPase assays were performed with full-length ClpX and the N-terminal deletion construct of ClpX (ΔNClpX). GFP-degron chimeras were used as substrates in these experiments. The deletion of the N-terminal domain does not substantially affect ATPase activity (which is ~1.2μM/min/μg, lower than full length ClpX ~1.4 μM/min/μg). While ClpX activity with the full-length enzyme showed a clear degron-dependent gradation-highest in the case of GFP_VAA_ (mimicking σ^D^/RsdA) and lowest for GFP_AAA_ (mimicking σ^L^/RslA), changes in specific activity of ΔNClpX were less pronounced (Figure 4D, Table 3-4). The unfoldase activity of ΔNClpX was also monitored for GFP-ssrA chimeras and compared with the activity of full-length ClpX. Consistent with previous observations, the unfoldase activity of ΔNClpX was less than ClpX while the degron dependence (GFP_VAA_ unfolded faster than GFP_GAA_ or GFP_AAA_) remained unaltered (Figure 4C, Table A1). To determine whether deletion of the N-terminal domain of ClpX affected the binding affinity with the GFP-degron substrate, we performed SPR experiments with the ΔNClpX construct. There was a two-fold decrease in the binding affinity of substrates to ΔNClpX when compared to full-length ClpX (Figure A4, Table A2). Despite the lower binding affinity for substrate proteins, the ΔNClpX construct retained degron-dependent gradation-highest in the presence of GFP_VAA_ and lowest for GFP_AAA_.

**Table 4:**
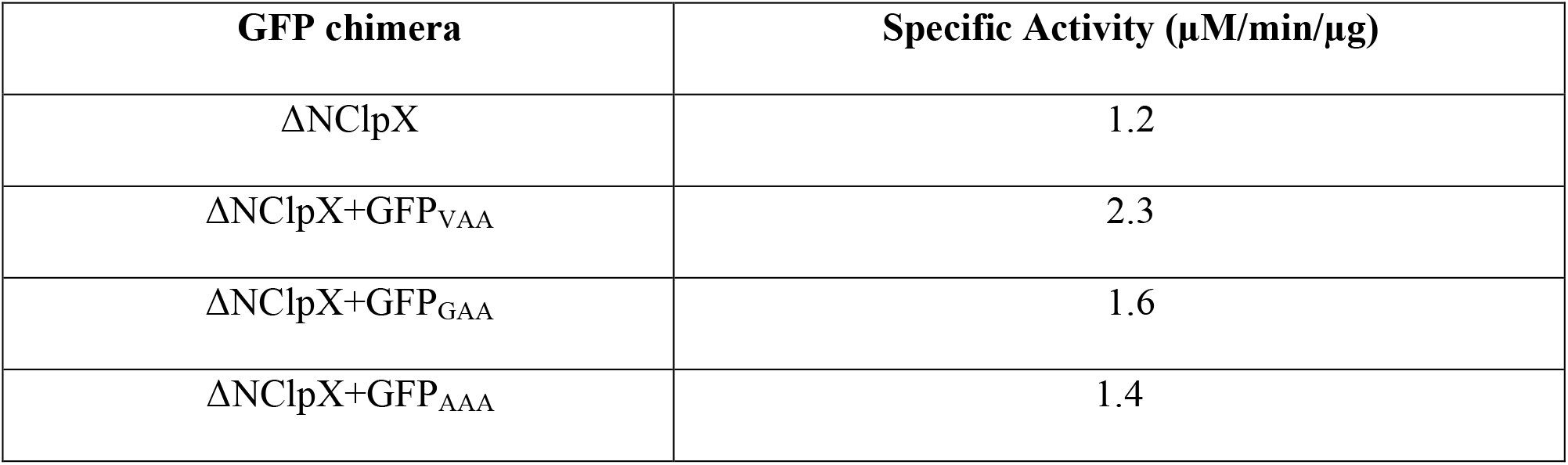
ATPase activity of ΔNClpX in the presence of GFP-chimeras.

ATP binding was shown to stabilise the ClpX hexamer (10). An interesting observation from these experiments is that ClpX substrate interactions can occur in the absence of nucleotides. Since this observation stands out from the *E. coli* ClpX system and that the possibility of the ΔNClpX construct being more unstable than the full-length ClpX persists, we experimentally evaluated the binding affinity of GFP_VAA_ for both ClpX and ΔNClpX in the presence of ATP (Figure A5). Although addition of ATP improved substrate binding for both ClpX and ΔNClpX constructs, ΔNClpX bound substrates with lower affinity when compared to full-length ClpX (Figure A5, Table A3). The N-terminal domain thus influences binding affinity and the residence time of substrate proteins on ClpX.

### ClpX links proteostasis with transcription

Under diverse stress conditions, proteolytic degradation of an anti-σ factor releases a free ECF σ factor to bind to the RNA polymerase and initiate transcription. In the case of the *E. coli* anti-σ factor RseA, apart from ClpX, other cellular proteases like ClpAP, Lon and FtsH were shown to mediate proteolytic degradation (7). *M. tuberculosis* has three Clp-unfoldases-ClpX, ClpC1 and ClpB. In the case of the cytosolic *M. tuberculosis* σ^E^/RseA complex, phosphorylated RseA was shown to be a target for proteolytic processing by the ClpC1P2P1 assembly (31). ClpB was shown to be primarily involved in the prevention of heat induced aggregation and refolding of denatured proteins (32-34). Homologues of the *E. coli* Lon and HslUV proteases are absent in *M. tuberculosis* (35,36). In the light of these observations, *M. tuberculosis* ClpC1 appeared to be the other likely unfoldase that could participate in anti-σ factor degradation. Experiments performed with freshly purified *M. tuberculosis* ClpXP2P1 and ClpC1P2P1 complexes reveal that σ^D^/RsdA is proteolysed specifically by the ClpXP2P1 and not by the ClpC1P2P1 complex (Figure 5A).

**Figure 5:**
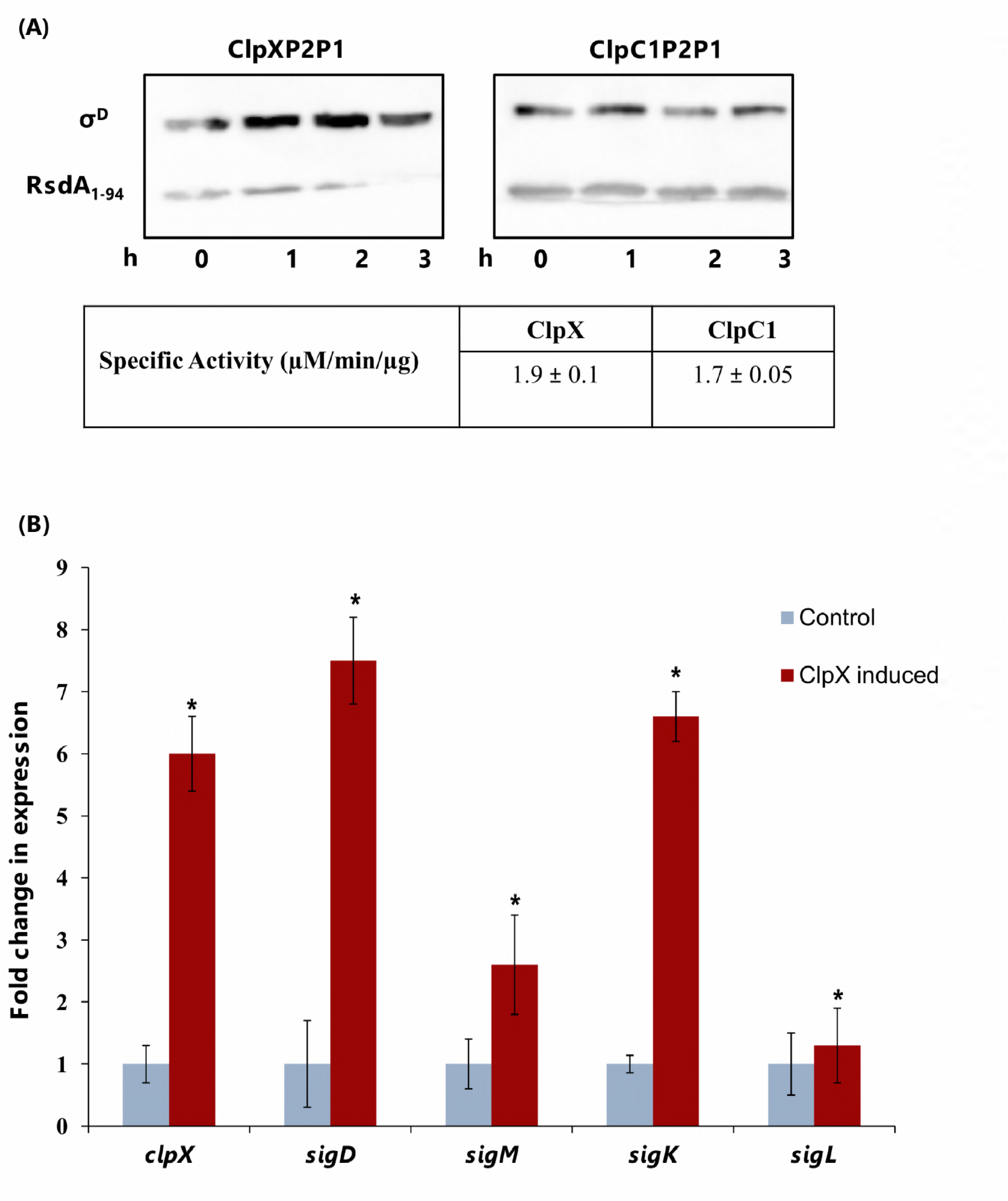
ClpX governs intracellular levels of membrane associated ECF σ factors. **(A) ClpX and not ClpC1 governs the degradation of an anti-σ factor with a ssrA-like degron.** For a comparison of the proteolysis of RsdA from the σ^D^/RsdA complex by the ClpXP2P1 and ClpC1P2P1 complexes, samples were analyzed for proteolysis at different time-points. This comparison suggests that RsdA is selectively proteolyzed by the ClpXP2P1 assembly. Targeted proteolysis was less pronounced with the ClpC1P2P1 assembly. The reaction mixture consisted of ClpX/ ClpC1: 0.3 μM, ClpP2P1: 0.8 μM, ATP (5 mM), σ^D^/RsdA substrate: 5.0 μM and an ATP-regeneration system (0.32 mg/ml Creatine kinase and 16 mM Creatine phosphate). Immunoblots were performed using antibodies raised against purified recombinant σ^D^ and RsdA. The ATPase activity of both unfoldases (ClpX and ClpC1) is similar. **(B) qPCR experiments reveal that the mRNA levels of all ECF σ factors increase upon ClpX induction.** The differences observed in the mRNA levels of *sigM* and *sigK* upon ClpX induction are broadly consistent with the relative degradation rates of their cognate anti-σ factors. Results shown here depict data from late stationary phase * indicates significance at p<0.05, ** indicates significance at p<0.005.

To evaluate if expression levels of *clpX* and the anti-σ factor genes were correlated, gene expression microarray datasets from different experimental conditions like hypoxia, stationary phase, oxidative stress and presence of Vitamin C were examined (compiled in Table 5). While σ^D^ maintains homeostasis in the late stationary phase of *M. tuberculosis* growth, genes in the σ^M^ regulon express in the stationary phase (37,38). The oxidative stress response involves both σ^K^ and σ^L^ as these ECF σ factors together respond to redox stress stimuli (27,28). The genes encoding for each *M. tuberculosis* σ and anti-σ factor pairs examined in this study lie in the same operon, and are positively regulated by the cognate σ factor. Despite multiple other factors that could influence gene expression, we note a correlation between the expression level of *clpX* and *anti-σ* factor genes in specific environmental conditions (Table 5). This finding was further evaluated in an experiment wherein the mRNA levels of the different σ factors were monitored upon *clpX* induction by quantitative real-time PCR (qPCR). The premise for this experiment was that increasing ClpX levels would result in more degradation of the target anti-σ factor thereby enhancing the cellular abundance of the corresponding free ECF σ’s. An increase in the intracellular levels of ECF σ’s, in turn, would result in upregulation of genes in the corresponding regulon. This hypothesis was examined in two experimental conditions in *M. tuberculosis* H37Rv-at the logarithmic phase and late stationary phase of growth. The stationary phase, in particular, was evaluated as RsdA (anti-σ^D^) shows the highest susceptibility for ClpX induced proteolysis *in vitro* and σ^D^ was shown to maintain homeostasis in the late stationary phase of *M. tuberculosis* growth (39). We note that the expression levels of σ^D^ are most upregulated upon ClpX induction in both experimental conditions (Figure 5B, A5). On the other hand, the mRNA levels of *sigL* were relatively unaffected upon *clpX* induction. Thus increased levels of ClpX directly influence the intracellular concentration of free σ factors. We note that under logarithmic growth phase (in which the anti-σ factor is less susceptible to RIP proteolysis) the effect is less pronounced (Figure A6, A7).

**Table 5:** Correlation between the expression of *clpX* and *anti-σ* factors*.

| Genes | Log phase | Stationary phase | Hypoxia | R.O.S | Vitamin C |
| --- | --- | --- | --- | --- | --- |
| *rsdA* | -0.59 (-1.5) | -0.16 (-0.4) | **1 (2.5)** | **0.96 (2.4)** | **0.99 (2.5)** |
| *rsmA* | **-0.95 (-2.4)** | -0.72 (-1.8) | 0.8 (2) | 0.32 (0.8) | -0.42 (-1) |
| *rskA* | **0.99 (2.5)** | 0.78 (1.9) | **0.93 (2.4)** | 0.38 (0.9) | 0 (0.08) |
| *rslA* | 0.53 (1.3) | **0.91 (2.3)** | 0.54 (1.4) | -0.27 (-0.7) | 0.49 (1.3) |

Abbreviations: *rsdA*: anti- $\sigma^D$ ; *rsmA*: anti- $\sigma^M$ ; *rskA*: anti- $\sigma^K$ ; *rslA*: anti- $\sigma^L$ .
Legend: Correlation coefficients with values $>0.9$ are in bold. The numbers in parenthesis indicate t-values. \*These correlations were compiled using microarray data (Geo accession nos. GSE16146, GSE101048 and GSE8786).

In an effort to evaluate down-stream effects of changes in σ factor levels, the mRNA levels of representative genes from the regulons of *sigD, sigM, sigK* and *sigL* were examined by qPCR in the late stationary phase (40-42). These experiments reveal that (i) The expression of all four ECF σ factors is upregulated upon ClpX induction and (ii) Upregulation of the cognate regulon is less clear (Figure 6A). Some aspects of the non-linear response upon ClpX induction can be rationalized, however. For example, while RskA and RslA share the same degron sequence (AAA), the expression of *sigK* (and rskA) is higher than *sigL* (and *rslA).* Another feature that could contribute to non-linearity is that the release of σ^K^ from the σ^K^/RskA complex is also governed by redox stimuli-σ^K^ is a redox sensor allowing dissociation of RskA under reducing conditions (28). It thus appears likely that differences in the expression of the σ^K^ regulon are likely to mimic steady state levels in stationary phase, low oxygen *M. tuberculosis* cultures (43). Another parameter that could significantly influence this experiment is that ClpX dependence is preceded by RIP-1 activity on the anti-σ factor in the Regulated Intramembrane Proteolysis (RIP) pathway. RIP-1 activity is also influenced by environmental stimuli (6). Taken together, these observations suggest that ClpX activity alters intracellular ECF σ factor levels in a degron-dependent manner thereby influencing the expression profile in *M. tuberculosis* (Figure 6B).

**Figure 6:**
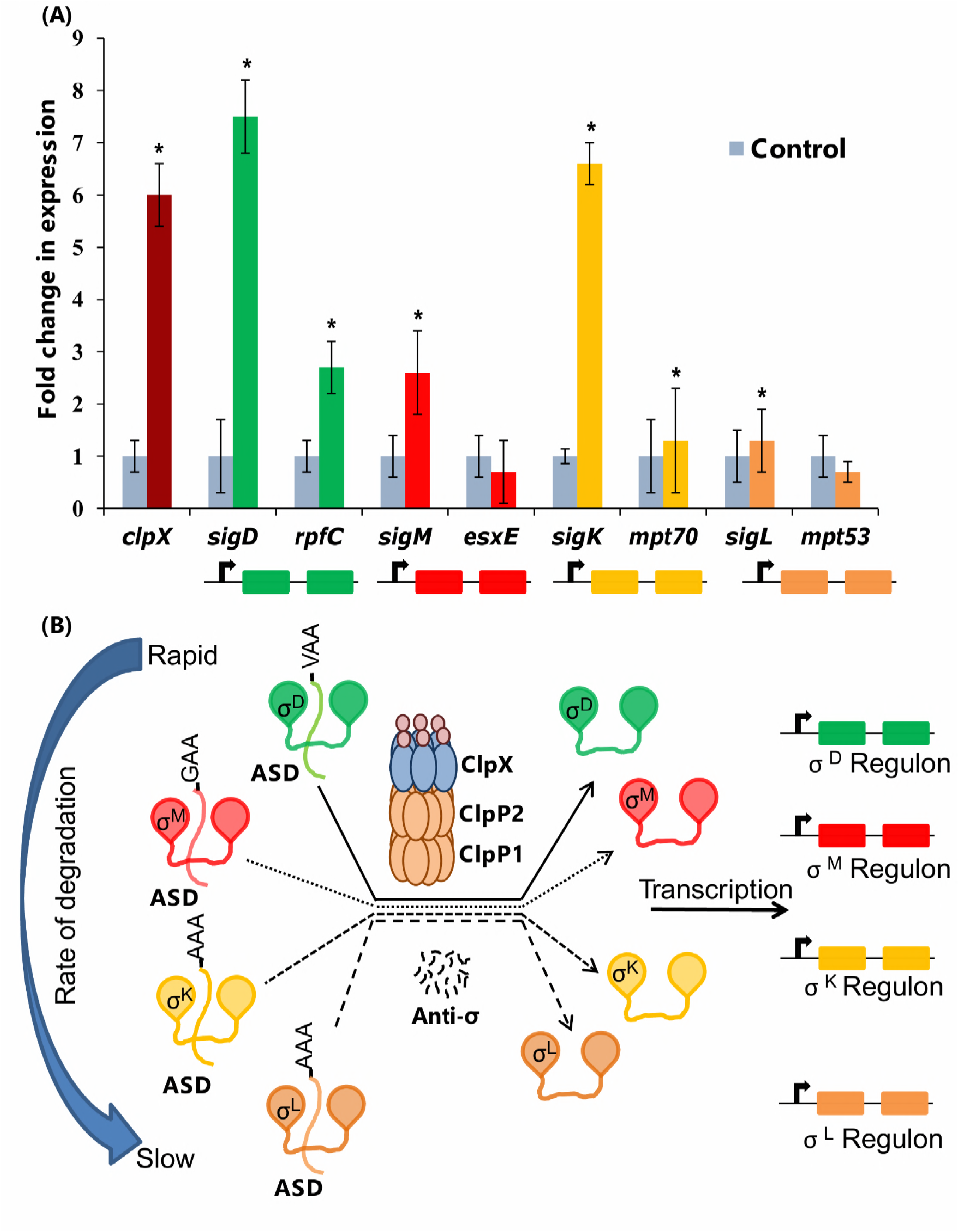
Mechanistic model for the gradient mechanism in the Regulated Intra-membrane Proteolysis (RIP) pathway. **(A) Fold change in mRNA levels of ECF σ factors and representative genes from their regulons.** While the expression of all ECF σ factors was upregulated upon ClpX induction, changes in the expression levels of representative genes *(rpfC, esxE, mpt70* and *mpt53)* in the target regulons is less pronounced (34,44-46). Multiple factors are likely to contribute to this non-linear response. Activation of σ^K^ and σ^L^ is also affected by redox stimuli (25,26). Other factors governing activation of σ^M^ remain to be determined. * indicates significance at p<0.05. **(B) A mechanistic model for degron-dependent variation in ClpX activity.** The rate of anti-σ factor degradation controls the cellular abundance of free ECF σ factors. Differences in the intracellular level of free σ factors alter the expression profile.

## Discussion

An intriguing feature in ECF σ factors, examined extensively in *E. coli* and *B. subtilis*, is that of overlapping regulons (44-47). This overlap in σ factor function ensures appropriate changes to the transcriptional profile - wherein multiple ECF σ’s are activated upon a stress signal (48). Another aspect is the apparent hierarchy amongst σ factors. Some σ factors *(M. tuberculosis* σ^C^ or σ^I^, for example) are under the control of σ^F^ which, in turn, is regulated by σ^M^ (49,50). Both these aspects depend on the cellular concentration of ECF σ’s which is largely regulated by a post-translational mechanism involving the release of free σ factor from an inactive σ/anti-σ complex (51). These release mechanisms either involve concerted conformational changes leading to the dissociation of the inactive σ/anti-σ complex or targeted proteolysis of the anti-σ to release the free ECF σ. Targeted proteolysis based on an accessible degron in a substrate effectively controls cellular protein levels. Indeed, this strategy is considered robust enough to be employed for optimizing microbial cell factories for diverse applications (52). It is in this context that the finding that the *M. tuberculosis* ClpX activity is modulated by degron composition becomes relevant.

In the Regulated Intramembrane Proteolysis (RIP) cascade, the signal transduction of environmental stress to the transcription mechanism has multiple temporal checkpoints-(i) the rate at which the extra-cytoplasmic receptor domain responds to the stress stimulus (ii) the rate of trans-membrane signal transduction involving the site-1 protease(s) and the site-2 protease Rip1 that acts on all membrane associated anti-σ’s (iii) the rate at which the anti-σ domain is selectively degraded by ClpX to release the free ECF σ to initiate transcription. In the *E. coli* RIP cascade, the proteolytic step initiated by the site-1 protease DegS is the rate limiting step (T_1/2_≤ 1 min) for RseA degradation, with the other two proteolytic events being at least three-fold faster (7). The dissociation of RseA from the membrane generates a cytosolic fragment with a degron (sequence ending in VAA) (8). The specific degradation of the cytosolic RseA by ClpXP is rapid (T_1/2_≤ 20 sec) and aided by an adaptor, SspB (7, 8). In the event ClpXP is weighed down by competing substrates, other cellular proteases can take over, albeit at a slower rate (T_1/2_≤ 1.6 min). Thus DegS activity on RseA is the rate-limiting step in the *E. coli* RIP pathway governing the cellular levels of σ^E^ (7). While *E. coli* RseA is potentially a substrate for multiple proteases like ClpA, HslUV, and Lon, ClpXP was demonstrated to be the major proteolytic complex in this process. It is worth noting in this context that *E. coli clpX* and *clpP* are not essential (53). On the other hand, *M. tuberculosis clpX, clpP1* and *clpP2* are essential genes (37,54).

The ability of *M. tuberculosis* ClpX to bind substrates in the absence of ATP suggested *M. tuberculosis* ClpX alternates between two different states, an observation similar to Hsp60 and Hsp70 chaperones (55). *E. coli* ClpX has also been shown to switch between two conformations- an ‘open state’ with lower binding affinity for substrates in the absence of nucleotides and a ‘closed state’ with higher affinity in response to ATP binding and hydrolysis (56). The finding that variations in substrate interaction kinetics elicited correlated changes in unfoldase activity suggested that concerted conformational changes could be a prominent feature of adaptor-independent ClpX activity. Indeed, in *E. coli,* conformational changes in AAA+ proteases were shown to provide a mechanism to correlate ATP hydrolysis with denaturation of target protein substrates (10). ATPase assays and SPR interaction studies performed with both wild-type ClpX and ΔNClpX suggest that the N-terminal domain plays a role in substrate recruitment by affecting both ATPase activity as well as substrate binding in ClpX. The correlation between interaction kinetics (monitored by Surface Plasmon Resonance) and ATPase activity was significantly dampened in ΔNClpX when compared to full-length ClpX. This observation that interaction kinetics obtained from SPR experiments are consistent with both ATPase activity and unfoldase assays (significant differences only in the presence of cognate substrates) suggests that non-specific interactions are unlikely. On the other hand, the observation that dissociation trajectories of substrates do not return to the baseline indicate a slow dissociation process-as aspect that has been reported earlier in the case of *E. coli* ClpX. A plausible rationale for this comes from the low frequency Normal Modes in the *M. tuberculosis* ClpX model that suggests flexible tethering of the NTD in ClpX could enable conformational changes for stronger substrate interactions (Figure A3). This finding is similar to *E. coli* ClpX wherein nucleotide-dependent movements of the NTD facilitate the entry of substrates inside the ClpX ring (57). Put together, these data are consistent with an induced fit model for ClpX that can be described as-

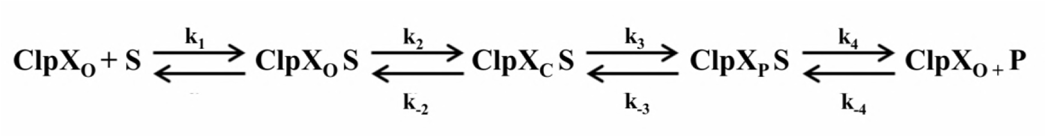

where k_1_ is the rate at which substrate binds, k_2_ and k_-2_ are the relative rates of conformational changes induced by substrate binding in the forward and backward directions, k_3_ is the rate of chemical reaction and k_4_ is the rate of product release. ‘o’ and ‘c’ stand for the open (before conformational change) and closed (after conformational change) states of the enzyme. The increase in ATPase activity upon *E. coli* ClpX interaction with a substrate was shown to precede a series of conformational changes in this enzyme (58,59). These transitions were seen to initiate conformational changes in the pore-1 (GYVG) and pore-2 (RKSENPSITRD) loops to accelerate ATP hydrolysis (58, 59). With a favourable amino acid at the ante-penultimate position, conformational changes engage the substrate and the enzyme switches from an open to closed state leading to increased ATPase activity and the consequent unfolding of the substrate. When k_-2_ becomes <<<< k_3_, the substrate is committed for the reaction after the conformational change. The observation that ATPase activity of ΔNClpX is less than full-length ClpX agrees well with the above model (Figure 4D, Table 3-4).

The finding that the last step in the RIP proteolytic cascade involving *M. tuberculosis* ClpX varies across substrates- fastest for RsdA and slowest for RslA- suggests significant differences from the *E. coli* model. In *E. coli,* the expression of the σ^E^ regulon in *clpX* and *sspB* null mutants is reduced-suggesting a correlation between proteolysis and transcription (8). The presence of multiple anti-σ’s in *M. tuberculosis* and degron-dependent variations in ClpX unfolding suggests a broader application of this link between targeted protein degradation and the expression profile. Previous reports on *sigA* transcript levels in *M. tuberculosis* H37Rv suggest that *sigA* expression is not altered at different phases of growth and exposure to stresses *in vitro* (44,60,61). However, upon ClpX overexpression, we observe differences in the levels of *sigA-* these are lower in the stationary phase than the logarithmic phase (Figure A8). σ^A^ modulates the expression of essential genes and virulence in *M. tuberculosis* (60). ClpX over-expression is thus likely to affect the ability of *M. tuberculosis* H37Rv to respond to stress. The degron-dependent differences in ClpX-anti-σ interactions and subsequent release of free ECF σ’s from the inactive complex is thus expected to lead to a measured change in transcription in response to a stress signal in *M. tuberculosis* (Figure 7A-B). An analysis using annotated σ/anti-σ pairs suggests that this feature is likely to be applicable in other bacteria (62). Indeed, a large proportion of ECF anti-σ factor sequences having one transmembrane helix (similar to the *M. tuberculosis* membrane associated anti-σ’s) have a ssrA-like degron (567 of 722 anti-σ factors) (Figure 1B). Put together, these data suggest that the *M. tuberculosis* ClpX function is more nuanced than a simple on/off switch in releasing ECF σ factors from an inactive σ/anti-σ complex. It appears likely that this variation in the unfoldase activity of ClpX is, in effect, a regulatory layer coordinating environmental stimuli to elicit calibrated changes in gene expression.

## Materials and Methods

### Cloning, expression and purification of recombinant proteins

*M. tuberculosis clpX, clpC1, clpP2* genes were cloned in the *E. coli* expression vector pET28a while *clpP1* was cloned in the MCSI of the pETDuet-1 vector (Novagen, Inc.). In case of the σ/anti-σ factor complex substrates, the full length σ factors (σ^D^, σ^K^, σ^L^ and σ^M^) were cloned in the multiple cloning site I (MCS-I), whereas the anti-σ factor constructs (ending at the ssrA-like motif at the C-terminal end), were cloned in the MCS-II of pETDuet-1 expression vector. GFP-ssrA from *E. coli* was obtained as a gift from Prof. Tania Baker’s laboratory. Mutants for the σ/anti-σ substrates and the GFP-ssrA mutants were prepared following standard Site-directed Mutagenesis (SDM) protocol (Table A4 lists the details of the constructs used in this study). The plasmids were transformed into a ClpP knockout strain of *E. coli* (obtained from Prof. Tania Baker’s laboratory). *E. coli* cultures were grown in Luria broth with appropriate antibiotic markers, to an optical density (O.D._600_) of 0.4-0.6 at 37°C, whereupon they were induced with 0.8mM isopropyl-β-D-1-thioglalactopyranoside (IPTG). Post induction, the cells were grown at a temperature of 18°C for 12-14 hours and harvested by centrifugation at 4500 rpm. The pellet for Clp-proteins was re-suspended and sonicated in lysis buffer (buffer L) containing 50mM Tris-HCl pH 7.6, 300mM NaCl, 100mM KCl, 1mM DTT, 10mM imidazole and 10% v/v glycerol, while the cell pellet for the σ/anti-σ factor complexes were re-suspended in buffer L devoid of DTT (except for the σ^L^/RslA complex). After sonication, the cell debris was separated from the crude cell lysate by centrifugation for 30 min at 15000 rpm. The cell-free lysate was then incubated with Ni^2^+-Nickel-nitrilotriacetic acid (NTA) affinity beads (Sigma-Aldrich, Inc.) for 1 hour at 4°C. The bound proteins were eluted by a gradient of imidazole concentration (50mM to 250mM) prepared in buffer L. The pure fractions were pooled, concentrated and loaded on to a PD-10 desalting column (GE Healthcare) and were desalted in buffer D (50mM HEPES-KOH pH 7.5, 25mM MgCl_2_, 100mM KCl, 0.1mM EDTA and 10% v/v glycerol) for Clp-proteins, and buffer S (50mM HEPES-KOH, pH 7.5, 100mM KCl and 10% v/v glycerol) for the substrate proteins.

### Surface Plasmon Resonance

Interaction studies were performed on a BIACORE 2000 instrument (Biacore, Uppsala, Sweden). ClpX was covalently immobilized on a CM5 sensor chip (Biacore) using standardized protocol in replicates. The SPR buffer (50mM HEPES, 200mM KCl with 10% Glycerol at pH 7.5) filtered through 0.45 micron membrane filters (Millipore) and degassed was used in these experiments. Experiments were carried out at 25°C. Carboxymethyl groups on the chip were activated by injecting freshly prepared Ethyl-3-(3-dimethylaminopropyl)-carbodiimide/ N-Hydroxysuccinimide (EDC/NHS: 1M each) mixture (1:1). ClpX diluted in 10mM Sodium acetate (pH 4.0) was then passed over the active surface till required immobilization was achieved. The un-reacted activated sites were blocked with 1M Ethanolamine. 50 μl (flow rate: 30μl/min) of each substrate at various concentrations were passed over the flow cells and allowed to dissociate for 200 seconds. The sensor surface was regenerated using multiple injections of 4M MgCl_2_ and/or 0.05-0.1% SDS whenever required. The reference subtracted response curves obtained for substrate binding to ClpX were evaluated using BIA evaluation software. The data obtained was fit to Langmuir 1:1 interaction model to obtain rates of association (k_a_) and dissociation (k_d_). Standard deviation across replicates was used to calculate the fitting error (Table A5, Figure A2) (63,64). The equilibrium dissociation constant (K_D_) is defined as the ratio of the dissociation rate constant (k_d_) and the association rate constant (k_a_).

### ATPase assay

ATPase assays were performed using malachite green to calculate the specific activity of ClpX. The assays were performed with 100nM ClpX and 5μM of substrate protein or 20μM of ssrA peptides in buffer containing 25 mM HEPES (pH 7.6), 200 mM KCl, 20 mM MgCl_2_, 10% glycerol. The reaction was initiated by the addition of 1mM ATP and was carried out at 30°C for 25mins. Malachite green dye buffer containing 0.045% malachite green, 4.2% ammonium molybdate and 1% Triton X-100 was added to the reaction mixture at suitable time points. After 1 min, 34% citric acid was added to the reaction mixture, mixed well and further incubated for 40mins for colour development. Absorbance was measured at 660nm. The inorganic phosphate released was calculated based on the absorbance standard curve established by H_3_PO_4_ standards.

### Unfoldase Assay

The catalytic unfolding of GFPssrA substrate by ClpX was monitored using an identical experimental protocol as in the case of *E. coli* ClpX (56). Unfolding of GFP_ssrA_ was monitored using a Varioskan plate reader with an excitation wavelength of 488nm and emission wavelength of 520nm. The reaction was monitored over a period of 30 minutes (Figure A9). The reaction mixture comprised of 1X Unfoldase buffer (25mM HEPES pH 7.5, 20mM MgCl2, 10% glycerol), 1X ATP regeneration system (Creatine Phosphate (16mM) and Creatine Kinase (0.32mg/ml), 200mM KCl, 100nM ClpX and 5uM of GFP_ssrA_ substrate.

### Molecular modelling and Normal Modes Analysis

The molecular model for *M. tuberculosis* ClpX was constructed using Modeller (65). The crystal structure of *E. coli* ClpX (3HWS) was used as template for comparative modelling. In this procedure, care was taken to ensure that the asymmetry observed in the ClpX hexamer was retained in the energy minimised model. Both the C and F chains of the obtained model had the closed conformation as observed in the *E. coli* ClpX crystal structure. Molecular graphics, energy minimisation of the model and analyses were performed with the UCSF Chimera package (66). Normal Modes Analysis was performed using Anisotropic Network Model Web Server 2.1 (67).

### RNA isolation and qPCR analysis

*M. tuberculosis* H37Rv transformed with pNit-3F vector and pNit-3F-ClpX were grown in the presence of 5 μM Isovaleronitrile (IVN) as inducer. 10 ml of bacterial cells at O.D._600_ ~1.0 were processed to extract RNA using the Trizol method. Briefly, cells were lysed in 1 ml Trizol using three cycles of 30s bead-beating with intermittent ice treatment for two minutes. The cell debris was removed from the lysate by centrifugation at 13,000 rpm for 10 minutes. The lysate was treated with 400 μl chloroform and centrifuged to separate the three phases. The top layer containing RNA was carefully extracted and the RNA was precipitated by addition of 1 ml isopropanol. The RNA pellet thus obtained was washed with 70% ethanol to remove excess salts. The dried RNA pellet was dissolved in 30μl RNase free water and kept overnight at 4°C to ensure complete dissolution. The RNA was quantified using NanoDrop (Thermo Scientific™ NanoDrop 2000c) and the sample was run on a 1% formaldehyde-agarose gel to estimate integrity. RNA sample was further processed by passing through RNeasy mini column (Qiagen). 1μg of purified RNA of each test and control sample was then treated with DNaseI (Thermo Scientific, Inc) to remove contaminating DNA and was used for cDNA synthesis employing one step-cDNA synthesis kit from Biorad (Biorad, Inc.). The cDNA synthesis was performed in 20μl reaction mixtures as per manufacturers’ protocol using 500ng of pure DNaseI treated RNA as template. Minus reverse transcriptase reaction was processed simultaneously as a control. For final qPCR reactions, the 20 μl of cDNA reaction mix was diluted to 100 μl and 2μl was utilised as template per reaction. Oligonucleotide sequences for qPCR were designed using ‘Primer 3’ software (Table A6 lists primers, Tm and the GC content of primers used). 16s rRNA amplicon was used as reference gene. Two-step SYBR green PCR reactions were performed in MasterCycler RealPlex4 (Eppendorf, Germany) machine as 10ul reactions in triplicates. The following conditions were used for amplification with SYBR green-10 minutes at 95°C followed by 40 cycles of 15 seconds at 95°C and 1 minute at 60°C. Melt curve analysis was done for each primer pair. Reverse transcriptase minus and plus cDNA samples were subjected to qPCR with 16s rRNA primers for analysis of gDNA contamination. For each individual gene analysed using qPCR, the C_q_ values were normalized with respect to C_q_ values of 16s rRNA amplicon. After normalization of C_q_ values, the fold change in the expression of various genes upon ClpX over-expression was calculated by 2^(−ΔΔCt)^ method as described previously. Relative quantification allowed us to relate the PCR signal of our target transcripts in ClpX over-expressed group to that of the Vector control (V.C.) group. The 2^(−ΔΔCt)^ method was used to analyze the relative changes in gene expression (68).

### Protease assays

The assays were performed in buffer containing 50 mM HEPES–KOH (pH 7.5), 150 mM KCl, 20 mM MgCl_2_ and 10% glycerol at 37°C. Clp proteases ClpP2 and ClpP1 (1μM each) were preincubated with 0.1mM Z-LL di-peptide for 30 minutes at 37°C. This step was followed by the pre-incubation of ClpC1 or ClpX (0.5 μM), ClpP2P1 (1μM), σ^D^/RsdA_VAA_ substrate (5.0 μM)/PyrB (1uM) substrate and an ATP-regeneration system (0.32 mg/ml Creatine kinase and 16 mM Creatine phosphate) for 10 minutes to allow formation of stable ClpXP2P1 or ClpC1P2P1 complexes. The reaction was initiated by the addition of 5mM ATP. Samples were removed after specific time intervals and a western blot was performed using anti-RsdA and anti-σ^D^ antibodies at 1:7000 dilution while ClpX, the Clp-proteases (ClpP2 and ClpP1) and PyrB were probed with anti-Histidine monoclonal antibodies (GE Healthcare) at 1:10,000 dilution. Immunoblots were developed with Luminata™ Forte Western HRP substrate for peroxidase-attached secondary antibodies.

### Expression levels and Correlation analysis

The expression levels of *clpX* and the four *anti-sigma* genes were obtained from previously published micro-array datasets (GSE16146, GSE101048 and GSE8786). Each dataset corresponds to different experimental conditions; Logarithmic phase (N=3), Stationary phase (N=3), Hypoxia (N=3), Oxidative Stress (Reactive Oxygen Species (R.O.S), N=3), Vitamin C (N=3). Pearson’s correlation coefficients and t values (Student’s t test) between a given *anti-σ* factor and *clpX* were calculated in all experimental conditions using the formula given below:

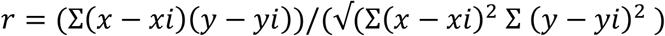

t=r/ (1-(0.92^2)) ^0.5 (r= correlation coefficient).

## Acknowledgements

We thank Ms. Shanta Sen and Mass spectrometry facility of National Institute of Immunology for mass spectrometry studies and Sneha Vishwanath, Himani Tandon for their help in computational studies. We thank Dr. Sess, Savitha N and Vandana and Ashish Deshmukh. We also thank Mrs. Sreelatha for her help with the SPR studies and Dr. Vinothkumar Kutti for his valuable comments on the manuscript.

## Author Contributions

ACJ, VKN and BG were involved in the design of this study. ACJ, PK, RKN, DSL were involved in data acquisition and analysis. ACJ, VKN and BG wrote the manuscript.

## References

1. Stallings CL, Glickman MS. 2010. Is *Mycobacterium tuberculosis* stressed out? A critical assessment of the genetic evidence. Microbes Infect., 12(14-15), 1091–1101.

2. Helmann JD. 2002. The extracytoplasmic function (ECF) sigma factors. Adv Microb Physiol., 46, 47–110.

3. Rodrigue S, Provvedi R, Jacques PE, Gaudreau L, Manganelli R. 2006. The sigma factors of Mycobacterium tuberculosis. FEMS Microbiol. Rev., 30, 926–941.

4. Sachdeva P, Misra R, Tyagi AK, Singh Y. 2009. βThe sigma factors of Mycobacterium tuberculosis: regulation of the regulators, FEBS J., 277, 605–626.

5. Gruber TM, Gross CA. 2003. Multiple sigma subunits and the partitioning of bacterial transcription space. Annu. Rev. Microbiol., 57, 441–466.

6. Heinrich J, Wiegert T. 2009. Regulated intramembrane proteolysis in the control of extracytoplasmic function sigma factors. Res. Microbiol., 160, 696–703.

7. Chaba R, Grigorova IL, Flynn JM, Baker TA, Gross CA. 2007. Design principles of the proteolytic cascade governing the σE-mediated envelope stress response in *Escherichia coli*: keys to graded, buffered, and rapid signal transduction. Genes Dev., 21, 124–136.

8. Flynn JM, Levchenko I, Sauer RT, Baker TA. 2004. Modulating substrate choice: the SspB adaptor delivers a regulator of the extracytoplasmic-stress response to the AAA+ protease ClpXP for degradation. Genes & Dev., 18, 2292–2301.

9. Flynn JM, Levchenko I, Seidel M, Wickner SH, Sauer RT, Baker TA. 2001. Overlapping recognition determinants within the ssrA degradation tag allow modulation of proteolysis. Proc Natl Acad Sci.USA, 98(19), 10584–10589.

10. Sauer RT, Baker TA. 2012. ClpXP, an ATP-powered unfolding and protein-degradation machine. Biochem Biophys.Acta., 1823(1), 15–28.

11. Mahmoud SA and Chien P. 2018, Regulated Proteolysis in Bacteria. Annu. Rev. Biochem. 87, 677–696.

12. Dougan DA, Mogk A, Zeth K, Turgay K, Bukau B. 2002. AAA+ proteins and substrate recognition, it all depends on their partner in crime. FEBS Lett., 529, 6–10.

13. Neher SB, Sauer RT, Baker TA. 2003. Distinct peptide signals in the UmuD and UmuD' subunits of UmuD/D' mediate tethering and substrate processing by the ClpXP protease. Proc Natl Acad Sci U S A., 100, 13219–13224.

14. Miller JM, Enemark EJ. 2016. Fundamental Characteristics of AAA+ Protein Family Structure and Function. Archaea, Vol.2016, Article ID 9294307, 12pages.

15. Wojtyra UA, Thibault G, Tuite A, Houry WA. 2003. The N-terminal Zinc Binding Domain of ClpX Is a Dimerization Domain That Modulates the Chaperone Function. J Biol Chem., 278(49), 48981–48990.

16. Donaldson LW, Wojtyra U, Houry WA. 2003. Solution Structure of the Dimeric Zinc Binding Domain of the Chaperone ClpX. J Biol Chem., 278(49), 48991–48996.

17. Glynn SE, Martin A, Nager AR, Baker TA, Sauer RT. 2009. Structures of Asymmetric ClpX Hexamers Reveal Nucleotide-Dependent Motions in a AAA+ Protein-Unfolding Machine. CELL., 139(4), 744–756.

18. Hanson PI, Whiteheart SW. 2005. AAA+ Proteins: Have Engine, Will Work. Nat Rev Mol Cell Biol., 6(7), 519–529.

19. Flynn JM, Neher SB, Kim YI, Sauer RT, Baker TA. 2003. Proteomic discovery of cellular substrates of the ClpXP protease reveals five classes of ClpX-recognition signals. Mol Cell., 11(3), 671–683.

20. Sklar JG, Makinoshima H, Schneider JS, Glickman MS. 2010. M. tuberculosis intramembrane protease Rip1 controls transcription through three anti-sigma factor substrates. Mol Microbiol., 77(3), 605–617.

21. Schneider JS, Sklar JG, Glickman MS. 2014. The Rip1 Protease of *Mycobacterium tuberculosis* Controls the SigD Regulon. J Bacteriol., 196(14), 2638–2645.

22. Alba BM, Leeds JA, Onufryk C, Lu CZ, Gross CA. 2002. DegS and YaeL participate sequentially in the cleavage of RseA to activate the sigma(E)-dependent extracytoplasmic stress response. Genes Dev. 16(16), 2156–68.

23. Kanehara K, Ito K, Akiyama Y. 2002. YaeL (EcfE) activates the sigma(E) pathway of stress response through a site-2 cleavage of anti-sigma(E), RseA. Genes Dev. 16(16), 2147–55.

24. Heinrich J, Hein K and Wiegert T 2009. Two proteolytic modules are involved in regulated intramembrane proteolysis of Bacillus subtilis RsiW. Mol. Microbiol., 74, 1412–1426.

25. Raju RM, Unnikrishnan M, Rubin DHF, Krishnamoorthy V, Kandror O, Akopian TN, Goldberg AL, Rubin EJ. 2012. Mycobacterium tuberculosis ClpP1 and ClpP2 Function Together in Protein Degradation and Are Required for Viability in Vitro and during Infection. PloS Pathog., 8(2), e1002511.

26. Jaiswal RK, Prabha TS, Manjeera G, Gopal B. 2013. Mycobacterium tuberculosis RsdA provides a conformational rationale for selective regulation of σ-factor activity by proteolysis. Nucleic Acids Res. 41(5), 3414–3423.

27. Thakur KG, Praveena T, Gopal B. 2010. Structural and Biochemical Bases for the Redox Sensitivity of *Mycobacterium tuberculosis* RslA. J. Mol.Biol., 397(5), 1199–1208.

28. Shukla J, Gupta R, Thakur KG, Gokhale R, Gopal B. 2014. Structural basis for the redox sensitivity of the *Mycobacterium tuberculosis* SigK-RskA σ-anti-σ complex. ActaCrystallogr D., 70, 1026–1036.

29. Thakur KG, Jaiswal RK, Shukla JK, Praveena T, Gopal B. 2010. Over-expression and purification strategies for recombinant multi-protein oligomers: A case study of *Mycobacterium tuberculosis* σ/anti-σ factor protein complexes. Prot. Exp. Purif., 74(2), 223–230.

30. Burton RE, Baker TA, Sauer RT. 2003. Energy-dependent degradation: Linkage between ClpX-catalyzed nucleotide hydrolysis and protein-substrate processing. Protein Sci., 12(5), 893–902.

31. Barik S, Sureka K, Mukherjee P, Basu J, Kundu M. 2010. RseA, the SigE specific anti-sigma factor of Mycobacterium tuberculosis, is inactivated by phosphorylation-dependent ClpC1P2 proteolysis. Mol. Microbiol., 75, 592–606.

32. Goloubinoff P, Mogk A, Zvi AP, Tomoyasu T, Bukau B. 1999. Sequential mechanism of solubilization and refolding of stable protein aggregates by a bichaperone network. Proc Natl Acad Sci USA., 96, 13732–13737.

33. Mogk A, Tomoyasu T, Goloubinoff P, Rudiger S, Roder D, Langen H, Bukau B. 1999. Identification of thermolabile *Escherichia coli* proteins: prevention and reversion of aggregation by DnaK and ClpB. EMBO J., 18, 6934–6949.

34. Motohashi K, Watanabe Y, Yohda M, Yoshida M. 1999. Heat-inactivated proteins are rescued by the DnaK·J-GrpE set and ClpB chaperones. Proc Natl Acad Sci USA., 96, 7184–7189.

35. Bajaj D, Batra JK. 2012. The C-terminus of ClpC1 of *Mycobacterium tuberculosis* is crucial for its oligomerization and function. PLoS One., 7, e51261.

36. Laederach J, Leodolter J, Warweg J, Weber-Ban E. 2014.. Chaperone-Proteases of Mycobacteria. In Houry, W. (ed.), The Molecular Chaperones Interaction Networks in Protein Folding and Degradation (pp. 445–481). Springer, New York.

37. Sassetti CM, Boyd DH, Rubin EJ. 2003. Genes required for mycobacterial growth defined by high density mutagenesis. Mol Microbiol., 48, 77–84.

38. Betts JC, Lukey PT, Robb LC, McAdam RA, Duncan K. 2002. Evaluation of a nutrient starvation model of *Mycobacterium tuberculosis* persistence by gene and protein expression profiling. Mol Microbiol. 43(3), 717–731.

39. Calamita H, Ko C, Tyagi S, Yoshimatsu T, Morrison N.E, Bishai W.R. 2004. The *Mycobacterium tuberculosis* SigD sigma factor controls the expression of ribosome-associated gene products in stationary phase and is required for full virulence. Cell Microbiol.; 7:233–244.

40. Raman S, Hazra R Dascher CC, Husson RN. 2004. Transcription Regulation by the *Mycobacterium tuberculosis* Alternative Sigma Factor SigD and Its Role in Virulence. J. Bacteriol., 186(99), 6605–6616.

41. Hahn M, Raman S, Anaya M, Husson RN. 2005. The *Mycobacterium tuberculosis* Extracytoplasmic-Function Sigma Factor SigL Regulates Polyketide Synthases and Secreted or Membrane Proteins and Is Required for Virulence. J. Bacteriol., 187(20), 7062–7071.

42. Veyrier F, Saσd-Salim B, Behr MA. 2008. Evolution of the Mycobacterial SigK Regulon. J. Bacteriol., 190 (6), 1891–1899.

43. Rosenkrands I, Slayden RA, Crawford J, Aagaard C, Barry CE, III, Andersen P. 2002. Hypoxic response of *Mycobacterium tuberculosis* studied by metabolic labeling and proteome analysis of cellular and extracellular proteins. J Bacteriol., 184:3485–3491.

44. Wade JT, Castro Roa D, Grainger DC., Hurd D, Busby SJ, Struhl K, Nudler E. 2006. Extensive functional overlap between sigma factors in *Escherichia coli*. Nat Struct Mol Biol., 13(9), 806–814.

45. Luo Y, Helmann JD. 2009. Extracytoplasmic function sigma factors with overlapping promoter specificity regulate sublancin production in Bacillus subtilis. J. Bacteriol., 191, 4951–4958.

46. Cao M, Helmann JD. 2002. Regulation of the Bacillus subtilis bcrC bacitracin resistance gene by two extracytoplasmic function sigma factors. J Bacteriol., 184(22), 6123–6129.

47. Huang X, Fredrick KL, Helmann JD. 1998. Promoter recognition by Bacillus subtilis sigmaW: autoregulation and partial overlap with the sigmaX regulon. J Bacteriol., 180(15), 3765–3770.

48. Manganelli R, Dubnau E, Tyagi S, Kramer FR and Smith I. 1999. Differential expression of 10 sigma factor genes in Mycobacterium tuberculosis. Mol. Microbiol., 31, 715–724.

49. Chauhan R, Ravi J, Datta P, Chen T, Schnappinger D, Bassler KE, Balazsi G, Gennaro ML. 2015. Reconstruction and topological characterization of the sigma factor regulatory network of Mycobacterium tuberculosis. Nat. Commun., 7, 11062.

50. Bervoets I, van Brempt M, van Nerom K, van Hove B, Maertens J, de Mey M, Charlier D. 2018. A sigma factor toolbox for orthogonal gene expression in *Escherichia coli*. Nucleic Acids Res., 46(4), 2133–2144.

51. Campbell EA, Westblade LF, Darst SA. 2008. Regulation of bacterial RNA polymerase σ factor activity: a structural perspective. Curr Opin Microbiol., 11, 121–127.

52. Landry BP, Stöckel J, Pakrasi HB. 2013. Use of Degradation Tags To Control Protein Levels in the Cyanobacterium *Synechocystis sp*. Strain PCC 6803. Appl Environ Microbiol., 79(8), 2833–2835.

53. Gottesman S. 1996. Proteases and their targets in *Escherichia coli*. Annu. Rev. Genet., 30, 465–506.

54. Ollinger J, O'Malley T, Kesicki EA, Odingo J, Parish T. 2012. Validation of the essential ClpP protease in *Mycobacterium tuberculosis* as a novel drug target. J. Bacteriol., 194, 663–668.

55. Bukau B, Horwich AL. The hsp70 and hsp60 chaperone machines. Cell. 1998;92:351–366.

56. Singh SK, Grimaud R, Hoskins JR, Wickner S, Maurizi MR. 2000. Unfolding and internalization of proteins by the ATP-dependent proteases ClpXP and ClpAP. Proc Natl Acad Sci U S A. 97:8898–8903.

57. Thibault G, Tsitrin Y, Davidson T, Gribun A, Houry WA. 2006. Large nucleotide-dependent movement of the N-terminal domain of the ClpX chaperone. EMBO J., 25:3367–3376.

58. Martin A, Baker TA, Sauer RT. 2008. Pore loops of the AAA+ ClpX machine grip substrates to drive translocation and unfolding. Nat Struct Mol Biol., 15(11), 1147–1151.

59. Siddiqui SM, Sauer RT, Baker TA. 2004. Role of the processing pore of the ClpX AAA+ ATPase in the recognition and engagement of specific protein substrates. Genes Dev., 18(4), 369–374.

60. Wu S, Howard ST, Lakey DL, Kipnis A, Samten B, Safi H, Gruppo V, Wizel B, Shams H, Basaraba RJ, Orme IM, Barnes PF. 2004. The principal sigma factor sigA mediates enhanced growth of *Mycobacterium tuberculosis* in vivo. Mol. Microbiol. 51, 1551–1562.

61. Hu YM, Coates ARM. 1999. Transcription of two sigma 70 homologue genes, sigA and sigB, in stationary-phase *Mycobacterium tuberculosis*. J Bacteriol. 181:8.

62. Staroń A, Sofia HJ, Dietrich S, Ulrich LE, Liesegang H, Mascher T. 2009. The third pillar of bacterial signal transduction: classification of the extracytoplasmic function (ECF) sigma factor protein family. Mol Microbiol., 74(3), 557–581.

63. Ortega J., Singh,S.K., Maurizi,M.R. and Steven,A.C. (2000) Visualization of substrate binding and translocation by the ATP-dependent protease. Mol. Cell, 6, 1515–1521.

64. Sugimoto S, Yamanaka K, Nishikori S, Miyagi A, Ando T, Ogura T (2010) AAA+ chaperone ClpX regulates dynamics of prokaryotic cytoskeletal protein FtsZ. J Biol Chem 285: 6648–6657.

65. Šali A, and Blundell TL. 1993. Comparative protein modelling by satisfaction of spatial restraints. J. Mol. Biol., 234, 779–815.

66. Pettersen EF, Goddard TD, Huang CC, Couch GS, Greenblatt DM, Meng EC, Ferrin TE. 2004. UCSF Chimera-a visualization system for exploratory research and analysis. J Comput Chem., 2(13), 1605–1612.

67. Eyal E, Lum G, Bahar I. 2015. The anisotropic Network Model web server at 2015 (ANM 2.0), Bioinformatics, 31, 1487–9.

68. Schmittgen TD, Livak KJ. 2008. Analyzing real-time PCR data by the comparative C(T)method. Nat Protoc., 3, 1101–1108.

70. Krogh A, Larsson B, von Heijne G, Sonnhammer EL. 2001. Predicting transmembrane protein topology with a hidden Markov model: application to complete genomes. J. Mol. Biol., 305, 567–580.

71. Bailey TL, Bodén M, Buske FA, Frith M, Grant CE, Clementi L, Ren J, Li WW, Noble WS. 2009. “MEME SUITE: tools for motif discovery and searching”. Nucleic Acids Res., 37, W202–W208.

72. Robert X, Gouet P. 2014. Deciphering key features in protein structures with the new ENDscript server. Nucl. Acids Res., 42(W1), W320–W324.

